# Dynamic changes in subplate and cortical plate microstructure at the onset of cortical folding in vivo

**DOI:** 10.1101/2023.10.16.562524

**Authors:** Siân Wilson, Daan Christiaens, Hyukjin Yun, Alena Uus, Lucilio Cordero-Grande, Vyacheslav Karolis, Anthony Price, Maria Deprez, Jacques-Donald Tournier, Mary Rutherford, Ellen Grant, Joseph V. Hajnal, A David Edwards, Tomoki Arichi, Jonathan O’Muircheartaigh, Kiho Im

## Abstract

Cortical gyrification takes place predominantly during the second to third trimester, alongside other fundamental developmental processes, such as the development of white matter connections, lamination of the cortex and formation of neural circuits. The mechanistic biology that drives the formation cortical folding patterns remains an open question in neuroscience. In our previous work, we modelled the in utero diffusion signal to quantify the maturation of microstructure in transient fetal compartments, identifying patterns of change in diffusion metrics that reflect critical neurobiological transitions occurring in the second to third trimester. In this work, we apply the same modelling approach to explore whether microstructural maturation of these compartments is correlated with the process of gyrification. We quantify the relationship between sulcal depth and tissue anisotropy within the cortical plate (CP) and underlying subplate (SP), key transient fetal compartments often implicated in mechanistic hypotheses about the onset of gyrification. Using in utero high angular resolution multi-shell diffusion-weighted imaging (HARDI) from the Developing Human Connectome Project (dHCP), our analysis reveals that the anisotropic, tissue component of the diffusion signal in the SP and CP decreases immediately prior to the formation of sulcal pits in the fetal brain. By back-projecting a map of folded brain regions onto the unfolded brain, we find evidence for cytoarchitectural differences between gyral and sulcal areas in the late second trimester, suggesting that regional variation in the microstructure of transient fetal compartments precedes, and thus may have a mechanistic function, in the onset of cortical folding in the developing human brain.

## Introduction

The gyrification of the human brain refers to the formation of grooves (sulci) and ridges (gyri) on the surface of the brain through the buckling and folding of the cortex during neurodevelopment. From an evolutionary perspective, this is thought to allow the expansion of cortical surface area within the confined space of the skull, optimizing the efficiency of brain circuitry by bringing interconnected cortical areas closer together, increasing the speed of information transmission (Fernández, Llinares-Benadero and Borrell, 2016). Alterations in the folding patterns of the cerebral cortex due to acquired injuries or congenital conditions can result in lifelong alterations in brain function, and have been implicated in the pathogenesis of intellectual disability, treatment-resistant epilepsy and various other cognitive or behavioral disorders (Nordahl *et al*., 2007; Barkovich *et al*., 2012; Subramanian, Calcagnotto and Paredes, 2020). Therefore, improving understanding of the mechanistic framework behind the early development of complex folding patterns during the human fetal period has key implications for improving the early diagnosis and clinical management of neurodevelopmental conditions.

After the bulk of neuronal migration has finished at the end of the second trimester, there is a rapid increase in the rate of cortical folding, resulting in the establishment of all the major sulcal landmarks of the adult brain by full term gestation (Chi, Dooling and Gilles, 1977; Yun *et al*., 2020). The deepest areas of the sulci, the sulcal pits, are the first to develop and are the most invariant across time, remaining spatially consistent even as other convolutions form (Lohmann, Von Cramon and Colchester, 2008; Im *et al*., 2010; Yun *et al*., 2020). The highly conserved nature of sulcal pits and primary folding patterns between individuals, compared to the secondary and tertiary sulci, have led to hypotheses that they are under closer genetic control than later developing sulci (Garel *et al*., 2001; Kostovic and Vasung, 2009; Im and Grant, 2019). By extension, the mechanisms governing the formation of cortical convolutions may differ pre- and post-natally, becoming more influenced by environmental factors at later stages of development. Although the timeline for the prenatal development of cortical surface features has been thoroughly described (Tallinen *et al*., 2016; Yun *et al*., 2020), the underlying in utero biology that drives cortical folding and determines individual variations in surface morphology is not well understood.

There are many models and hypotheses about the biological mechanisms that drive cortical gyrification (Van Essen, 1997, 2020; Lefèvre and Mangin, 2010; Xu *et al*., 2010; Tallinen *et al*., 2014, 2016), which seek to explain how the fundamental maturational processes of the second to third trimester interact dynamically to initiate the buckling of the cortex. These processes include the expansion of cortical surface area (Finlay and Darlington, 1995; Mota and Herculano-Houzel, 2015), dissipation of the subplate (SP) compartment (Rana *et al*., 2019), ingrowth of thalamocortical axons and white matter axonal tension (Van Essen, 1997, 2020), neuronal proliferation (Kriegstein, Noctor and Martínez-Cerdeño, 2006), regionally variant neuronal differentiation (Wang *et al*., 2017; Llinares-Benadero and Borrell, 2019), and differential expansion of cortical layers (Richman *et al*., 1975). Common to all of these hypotheses are the developmental processes occurring within the transient compartments of the fetal brain, the cortical plate (CP) and SP (Kostovic, 2002; Kostovic and Judas, 2007). In this work, we test the hypothesis that there is an association between microstructural development in the SP and CP and the onset of cortical folding, offering insight into the mechanism of cortical gyrification.

Diffusion MRI can be used to quantify structural maturation of the brain and can provide detailed information about intracellular and extracellular water diffusion over the second to third trimester (Takahashi *et al*., 2012; Huang and Vasung, 2014; Xu *et al*., 2014; Jaimes *et al*., 2020; Wilson *et al*., 2021, 2023). Importantly, modelling the diffusion signal allows a non-invasive yet refined interpretation of the dynamic changes seen in image contrast in terms of the underlying cellular biology or microstructure. This has been demonstrated in rodents (Sizonenko *et al*., 2007; Aggarwal *et al*., 2015), primates (Wang *et al*., 2017), human fetuses (Takahashi *et al*., 2012; Xu *et al*., 2014), and preterm infants (McKinstry, 2002; Ball *et al*., 2012; Batalle *et al*., 2019).

In our previous study (Wilson *et al*., 2023), we examined 112 brain MRI datasets from fetuses aged between 24 and 36 gestational weeks (GW) acquired as part of the developing Human Connectome Project (dHCP: https://nda.nih.gov/edit_collection.html?id=3955), using surface reconstructions of T2-weighted volumes and high angular resolution multi-shell diffusion-weighted imaging (HARDI) data. We initially describe the formation of sulci using sulcal depth, and then quantify how the microstructure of the underlying SP and CP layers changes with gestational age (GA) by modelling the diffusion MR signal with multi-shell multi-tissue constrained spherical deconvolution (MSMT-CSD). MSMT-CSD represents an optimized approach for the comparatively low signal to noise ratio and relatively high overall free water content in the developing brain (Pietsch *et al*., 2019; Wilson *et al*., 2021, 2023). We quantified how the tissue fraction, a measure of the proportion of anisotropic microstructure in a voxel, varies between transient fetal compartments and across gestational age. We characterised maturational trajectories that are complementary to histological understanding about the neurobiological processes that occur in these zones during the fetal period (Wilson *et al.,* 2023).

Projecting tissue fraction to the fetal surface, we quantify the relationship between emerging cortical folds and the microstructural properties of the underlying tissue layers, including both CP and SP. We correlate these metrics at the individual subject level within local patches across the surface, and find an inverse relationship between the anistropic tissue fraction and sulcal depth. We then project a sulcal depth map of folded brain regions onto the unfolded brain, to test the hypothesis that dynamic microstructural changes in the SP and CP precede the onset of cortical folding. We identify lower tissue fraction in regions which subsequently mature into sulcal fundi, prior to the formation of cortical folds during the second trimester. This result offers mechanistic insight into the microstructural changes that accompany the emergence of cortical folds, and the potential role of transient fetal compartments in this process.

## Results

### Cohort

For this study, we used a subset of the dHCP fetal cohort (www.developingconnectome.org, Edwards et al., 2024) as described previously (Wilson et al., 2023), consisting of 112 brain T2-weighted and Diffusion Weighted Imaging (DWI) datasets, age 24 to 36 weeks Gestational Age (GA) (Supplementary Figure 1), with an even ratio of male:female fetuses. The difference in cohort size to the original work reflects 28 subjects (all ≤ 24 weeks GA) that were discarded due to failed surface reconstruction and/or surface registration at younger gestational ages due to the absence of sulcal landmarks.

### Concomittant microstructural maturation and gyrification of the fetal brain over the second to third trimester

We used the CP segmentation of the T2w images to reconstruct the surface (Figure 1), then quantified the sulcal depth at each vertex in each subject, to chart the development of sulci over the second to third trimester (Figure 2a). Changes in sulcal depth at each primary sulci were best fitted by sigmoidal curves over this developmental window, following the expected pattern and timing of sulcal development (Yun *et al*., 2020) (Supplementary Figure 2).

**Figure 1.**
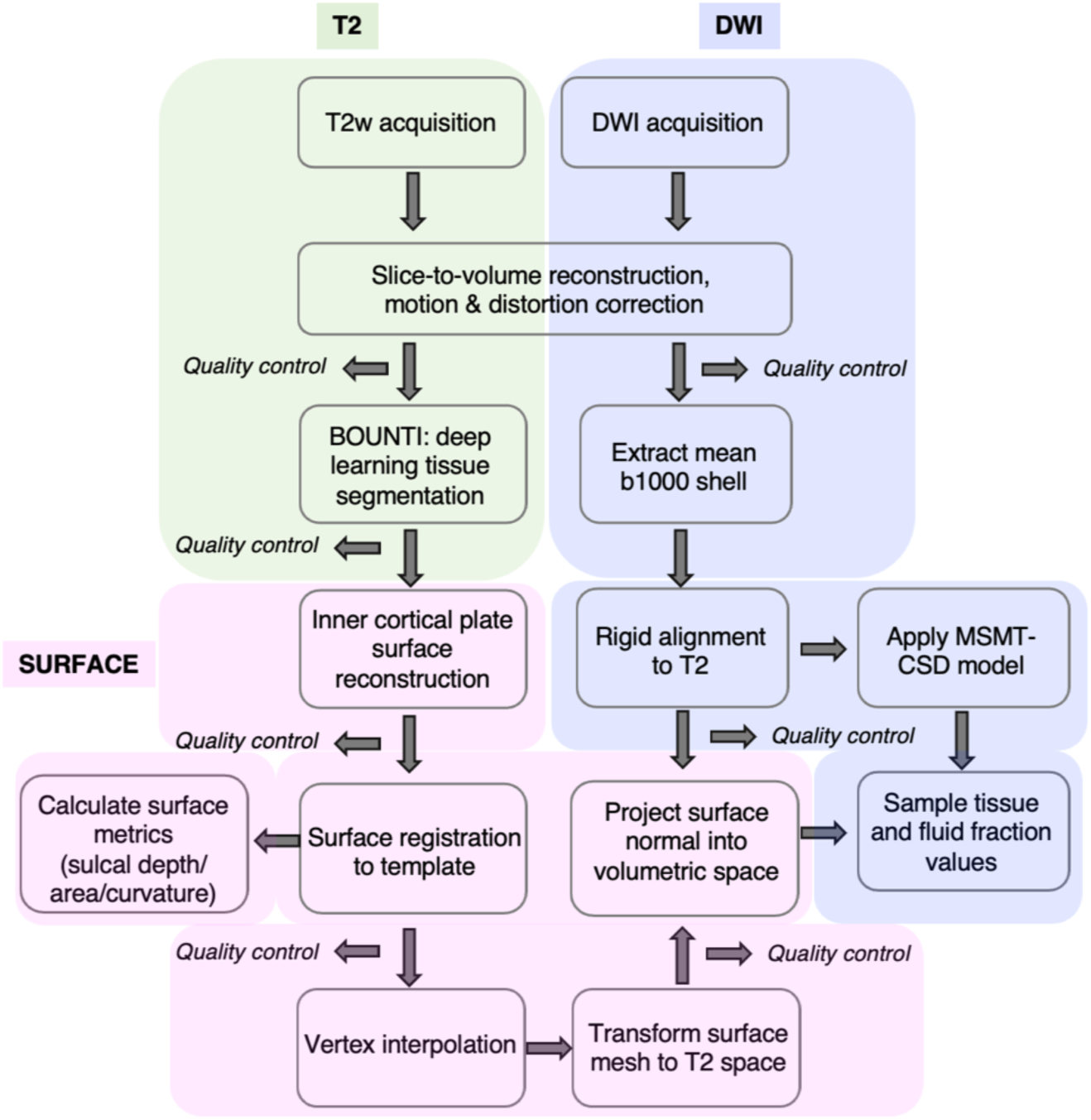
Imaging acquisition, preprocessing and analysis framework to integrate fetal T2w and DWI imaging for the investigation of microstructure on the fetal brain surface.

**Figure 2.**
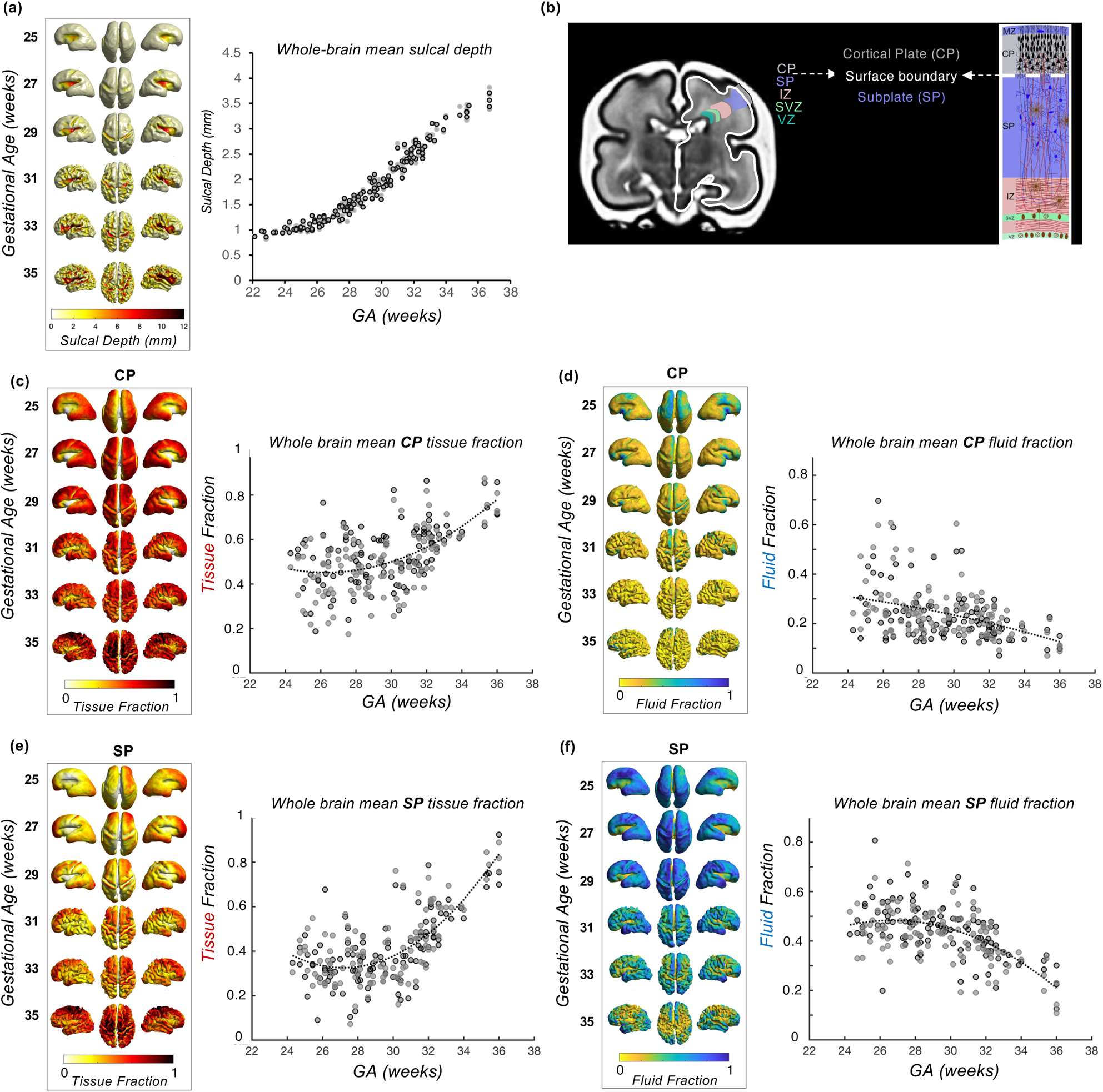
(a) Mean sulcal depth values across the surface at biweekly intervals and whole brain sulcal depth for each subject, plotted against GA in weeks (b) Illustration of the cortical plate (grey) and subplate (purple) compartments, separated by the surface boundary (white), and diagram of cytoarchitecture of fetal layers (adapted from Kostović and Judaš, 2015) (c) Biweekly mean CP tissue fraction and (d) fluid fraction for each surface vertex, projected to the surface and charted against GA (e) Biweekly mean SP tissue fraction and (f) fluid fraction for each surface vertex, projected to the surface and plotted against GA (grey = left, black = right), best fit with a 2^nd^ order polynomial curve.

As described in previous work (Wilson *et al.,* 2023), we quantify variations in microstructure using an advanced, data-driven modelling approach, MSMT-CSD. This is an alternative to the traditional diffusion tensor imaging model, which is confounded by its inability to successfully account for crossing fiber populations (Alexander *et al*., 2001; Alexander, Barker and Arridge, 2002). This limitation is particularly relevant in the fetal brain, which is characterised by incoherent immature axonal bundles, transient cellular structures, and more challenges due to increased partial volume effects inherent to imaging smaller brains (Jeurissen, 2013). Therefore, to adapt to the unique fetal imaging context, we applied a two-compartment version of MSMT-CSD to deconvolve the HARDI signal into the anisotropic, “white-matter-like”, tissue signal and the isotropic, CSF, “fluid” signal. This produced ‘tissue fraction’ and ‘fluid fraction’ maps, conveying the contributions from each component to the overall diffusion signal, which have been analyzed and published in our previous studies (Pietsch *et al.,* 2019, Wilson *et al.,* 2021, 2023).

We aligned the surfaces, T2 and DWI volumes for each subject, and used a ±2mm ribbon around the inner CP segmentation boundary to obtain diffusion metrics of the SP (inside boundary) and CP (outside boundary) at each vertex (Figure 2b). The 2 mm distance was chosen to match the resolution of the diffusion imaging (2 mm isotropic). To complement our previous findings (Wilson *et al.,* 2023), we summarise the maturational trajectory of the tissue and fluid fraction across the second to third trimester, taking the mean value for each metric in the CP and SP and plotted against GA (Figure 2c-f). We observe increasing tissue fraction with GA and decreasing fluid fraction in the CP and SP (Figure 2f), with a higher rate of change in the SP compared to the CP. This result was congruent with the variations in microstructure between these fetal compartments described in our previous work (Wilson et al., 2023).

Projecting these metrics to the surface highlighted regional variations in CP and SP microstructure across the brain, and as a function of GA (Figure 2c-f). In younger fetuses, we observe lower tissue fraction in the central sulcus and insular cortex, and higher tissue fraction in frontal and temporal lobes. Whereas at older GAs, the medial frontal and temporal pole have the highest tissue fraction. Fluid fraction generally showed a reciprocal pattern, except in the insula region, which also had relatively low fluid fraction values compared to other regions across GA.

### Tissue fraction decreases as a function of sulcal depth, independent of age

To test the hypothesis that microstructural maturation accompanies the formation of cortical folds, we analysed the coupling between cortical folding and diffusion metrics of the CP and SP, by correlating tissue fraction values against sulcal depth within local areas or ‘patches’ in individual subjects. Patches were defined by finding the neighbours around each vertex up to 5 degrees of separation (to create patches with a median of ∼100 vertices) (Figure 3a, see Methods). We then calculated the correlation between CP/SP tissue fraction values and sulcal depth between the vertices in each patch.

**Figure 3.**
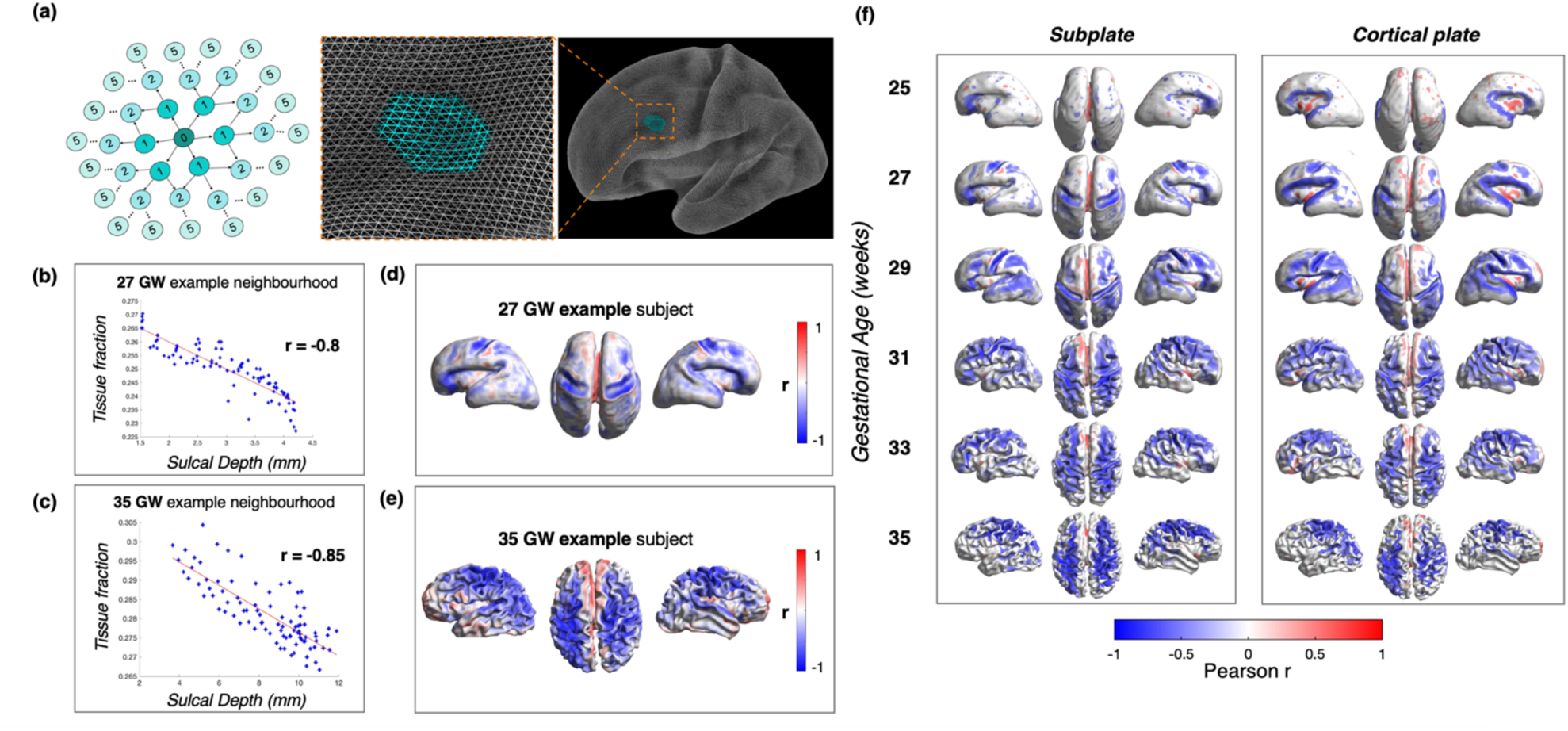
Within-subject tissue fraction vs. sulcal depth neighbourhood analysis (a) Schematic to demonstrate the neighbourhood concept for local patch analysis on the surface, and the degrees of separation between the central vertex and surrounding neighbours. (b) Example of 27 GW subject, and how ‘r’ value is derived, correlating sulcal depth and tissue fraction in a local patch in the central sulcus area (c) Example of neighbourhood in central sulcus at 35 GW. (d,e) Example subjects at whole brain level, with r values projected to the surface, colourbar represents strength of correlation (magnitude of r Pearson value). (f) Mean Pearson r for subjects grouped into biweekly bins, representing correlation between tissue fraction and sulcal depth in CP and SP across the whole brain (only significant values after FDR correction are shown, q = 0.05).

When examining individual subjects, we found many patches with a negative correlation between both CP and SP tissue fraction and sulcal depth, which corresponded to developing sulcal areas, such as the central sulcus at 27 GW (Figure 3b,d). This relationship was not only present at the onset of gyrification in younger subjects with early emerging sulci, but also in older subjects at the later stages of the third trimester as the cortical folds become deeper and more convoluted (Figure 3c,e). This result implies that at a global level, tissue fraction progressively decreases in both compartments down the sulcal bank into the sulcal fundus regions. To examine how prevalent this relationship was across the whole brain, we projected the Pearson r value at each vertex onto the surface (Figure 3d,e).

As the pattern of negative correlations across the surface was consistent between subjects of the same age, subjects were then grouped into biweekly bins, for which the mean Pearson’s correlation coefficient was calculated at each vertex (Figure 3f). FDR correction for multiple comparisons was applied across vertices, and then significant ‘r’ values (q < 0.05) were projected to the surface for each biweekly bin. We observed that this association was present in developing sulcal areas at every biweekly GA group (Figure 3f). Of note, the association was the most prominent in brain regions where sulci were forming, for example at 25 GW when significant neighbourhood associations were only seen in the sylvian fissure with progression then seen at 27 GW in the central sulcus, and by 35 GW in most primary sulci (Figure 3f).

### Tissue fraction minima are present in the unfolded brain, in regions that become sulci in the third trimester

To investigate whether changes in microstructure precede the emergence of cortical convolutions, we exploited the conserved spatial configuration of primary cortical folds between individuals, and back-projected a map of folded brain regions (sulcal depth) from the oldest subjects onto the unfolded brain in the late second trimester (Figure 4). We then used the same neighbourhood patches to calculate the correlation between mature sulcal depth values (35 - 37 GW) and younger tissue fraction values (25 – 30 GW) in an ‘age-mismatched’ analysis. This age-mismatched analysis (Figure 4) highlighted microstructural variations across sulcal areas in subjects age 25 – 30 GW prior to presence of particular folds, highlighting brain regions where tissue fraction in unfolded regions in younger subjects correlated with future sulcal depth values (Figure 4a,b).

**Figure 4.**
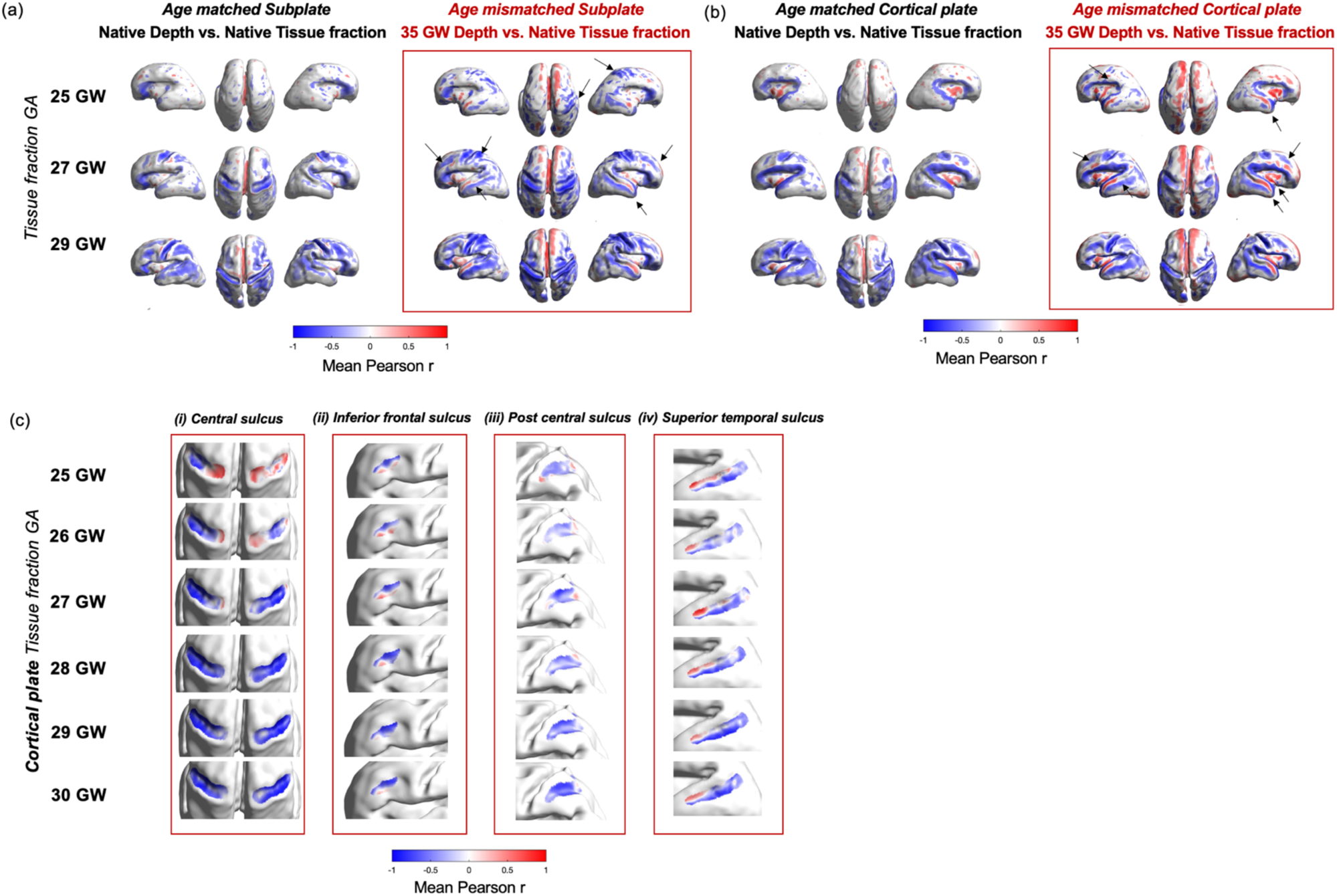
Age-mismatched analysis highlights brain regions where changes in microstructure precede the formation of sulcal folds, in the (a) SP and (b) CP. Red boxes are whole-brain correlation coefficient values for the relationship between sulcal depth at 35 GW and biweekly mean tissue fraction at 25, 27 and 29 GW. Arrows indicate areas of strong change between age matched and mismatched analysis, including the inferior frontal sulcus, superior temporal sulcus, post-central sulcus and insula. (c) Close-up, of correlation coefficient describing relationship between tissue fraction at 25 – 30 GW and sulcal depth at 35 GW in specific primary sulci, including (i) Central Sulcus, (ii) Inferior Frontal Sulcus (iii) Post central sulcus and (iv) Superior temporal sulcus.

When comparing the age-matched vs. age-mismatched maps (Figure 4a,b), we observed more regions with significant coupling in certain major sulcal landmarks in the age-mismatched scenario, including the inferior frontal sulcus, superior temporal sulcus and post-central sulcus (indicated with arrows). To explore this relationship in more detail, we performed specific analyses within sulcal regions of interest (ROIs) at each gestational week (Figure 4c). In the inferior temporal sulcus, post central sulcus and superior temporal sulcus, we observed strong correlation values between r = −0.56 and r = −0.97, which were consistent both spatially across the sulcus and temporally from 25 to 30 GW (Figure 4c(ii),(iii),(iv)). We also observe certain brain regions with positive correlations, particularly in the interhemispheric fissure and insula. Asymmetries between left and right hemispheres were noted in the central sulcus at early GAs, particularly in the youngest subjects, likely due to large inter-subject variability.

## Discussion

In this study, we characterise the microstructure of transient compartments in the fetal brain at the onset of the cortical folding process in the human brain in vivo. Specialised acquisition and image reconstruction techniques were designed to overcome the unique challenges associated with imaging the in utero environment, such as artefacts due to fetal head motion and geometric distortion due to magnetic inhomogeneities (Christiaens *et al*., 2021). These methods significantly enhanced data quality, enabling more reliable and accurate imaging analysis. With the unique dHCP fetal multi-shell HARDI dataset, it was possible to explore the relationship between the developing microstructure of the SP and CP, and the macrostructural appearance of sulci and gyri. We found regionally heterogenous patterns of cortical microstructural and macrostructural development, and a robust inverse relationship between tissue fraction and sulcal depth in specific emerging sulcal regions across the second half of in utero development. We then established that there is an inverse relationship between tissue fraction and ‘future’ sulcal depth across numerous key cortical areas, suggesting that the dynamic processes influencing microstructural properties in these compartments might be driving the emergence of cortical folds.

### Tissue fraction maturation reflects known developmental cortical processes in the cortical plate and subplate

Although it is known that fundamental developmental processes in the SP and CP increase microstructural maturity over the same timeframe that the primary and secondary cortical folds emerge (McKinstry, 2002; Ball *et al*., 2013; Huang and Vasung, 2014; Batalle *et al*., 2019), the precise relationship between them has not been established. This is particularly challenging as there are a multitude of biological processes occurring simultaneously in the CP and SP in the third trimester which create a complex microstructural environment. These include the disappearance of radial glial scaffolding, appearance of immature oligodendrocytes, dendritic arborization and axonal branching.

To complement our previous work (Wilson *et al.,* 2023), we used the two-compartment MSMT-CSD decomposition of the diffusion MRI, exploiting the multi-shell data and b-value dependency of different brain tissue types (Jeurissen *et al*., 2014). For subjects of the same GA, we observe high inter-subject correspondence in the spatial pattern of diffusion metrics across the surface. Similarly, we find subjects of the same age showed the same spatial distribution of significant ‘r’ values and identify a clear inverse relationship between tissue fraction and sulcal depth, localised to specific developing sulci. The inter-subject correspondence suggests this modelling and analysis approach offers sensitivity to developmental changes in signal properties, while also offering interpretable parameters with respect to biological tissue maturation (Pietsch *et al.,* 2019, Wilson *et al.,* 2023). The tissue fraction measure describes the proportion of anisotropic, coherently aligned microstructure within a voxel (Pietsch *et al.,* 2019; Wilson *et al.,* 2021), which increases with GA over the third trimester when an average is taken in the CP and SP layers. The maturation of this signal property fits with previous histological descriptions of maturing neuronal morphology, increases in cellular and organelle density, and reductions in extracellular water as dendritic trees develop and thalamocortical fibres establish synapses in the CP and SP (Kostovic and Judas, 2006; Sizonenko *et al*., 2007; Dean *et al*., 2013). Interestingly, we observed continuous tissue fraction increases in the CP across our study period; whereas in the SP, a rapid increase was not seen until after 30 GW (Figure 3). This developmental transition point within the subplate was also noted in previous studies (Wilson *et al*., 2021, 2023; Chen *et al*., 2022; Calixto *et al*., 2024) and in histological examinations showing the commencement of premyelination at approximately 30 GW, when immature oligodendrocytes begin extending their processes (Back *et al*., 2002; Kostovic and Vasung, 2009). Overall, these results highlight how the application of more complex data-driven modelling approaches to exploit the information in the multi-shell diffusion data can enable improved characterisation of the underlying biological complexity, which was not previously possible with in utero single-shell diffusion imaging.

### Tissue fraction values and FA are complementary but sensitive to different underlying biology

Previous in vivo and ex vivo work has characterised the developing microstructure of the cortex using DTI (McKinstry, 2002; Dudink *et al*., 2010; Ball *et al*., 2013; Vinall *et al*., 2013; Batalle *et al*., 2019). To place the novel MSMT-CSD derived metrics in the context of this previous work, we also charted FA changes in the CP and SP to compare the maturational trends (Supplementary Figure 3). Our findings are consistent with ex utero observations from studies across species, in preterm infants (Dudink *et al*., 2010; Batalle *et al*., 2019; Ouyang *et al*., 2019), non-human primates (Kostovic and Rakic, 1990; Kroenke *et al*., 2007) and rats (Huang *et al*., 2008), identifying a decrease in whole-brain FA in both the CP and SP, with a higher rate of change in the CP. This reflects a loss of radial coherence in the CP, the disassembly of radial scaffolding in the SP (as neuronal migration finishes at the end of the second trimester) and the isotropic restriction of water movement as dendritic arborization occurs (Sizonenko *et al*., 2007; Dean *et al*., 2013; Wang *et al*., 2017). We also observe decreases in MD, which likely relate to increases in synaptic, cellular and organelle density during this period (Sizonenko *et al*., 2007; Dean *et al*., 2013). These trends in DTI metrics have also been observed in preterm infants over a comparable timeframe in early development (McKinstry, 2002; Ball *et al*., 2013; Vinall *et al*., 2013; Batalle *et al*., 2019) and in post-mortem tissue from other gyrencephalic mammals such as ferrets and macaques (Kroenke *et al*., 2007, 2009; Yu *et al*., 2016). Crucially, the difference in the maturational trends of cortical tissue fraction and FA with GA suggests that although these metrics can be considered complementary for describing anisotropic, coherently aligned tissue-like signal, the models diverge due to their different sensitivities to specific components of the diffusion signal and thus capture distinct developmental process during the second to third trimester.

### Sulcal fundi are associated with a local decrease in tissue fraction, recapitulating adult cytoarchitectural organisation in utero

To investigate if the heterogeneous microstructural maturation in different cortical regions plays a mechanistic role in cortical folding, we quantified the relationship or ‘coupling’ between SP/CP microstructure and sulcal depth within local neighbourhoods or ‘patches’. We found a negative correlation in many developing sulcal areas at the individual subject level, which was spatially consistent between subjects of the same age. We observe these local reductions in tissue fraction in the deepest parts of sulci at the onset of cortical folding (in the sylvian fissure at 25 GW and in the central sulcus at 27 GW). Importantly, this trend emerges across the cortex as more regions fold, persisting throughout the cohort up to 35 GW as sulci deepen and mature. The higher tissue fractions we observe in gyral crowns are supported by previous studies across multiple gyrencephalic species, identifying higher myelinated fibre density, greater vertical alignment of pyramidal neurons, and longer more elaborate dendritic trees in gyral crowns than in sulcal pits (Nie *et al*., 2012; Mortazavi *et al*., 2017; Llinares-Benadero and Borrell, 2019). Our results are also consistent with ex vivo histological observations of a greater density of fiber bundles and persistence of radial pathways in developing gyral areas compared to sulcal areas (Takahashi *et al*., 2012). Overall, our analysis reveals that the neuroanatomical differences between gyral and sulcal areas observed in adulthood are already present at the formation of the folds in utero, suggesting this conformational difference in microstructure has mechanistic importance (see infographic summary in Figure 5).

**Figure 5.**
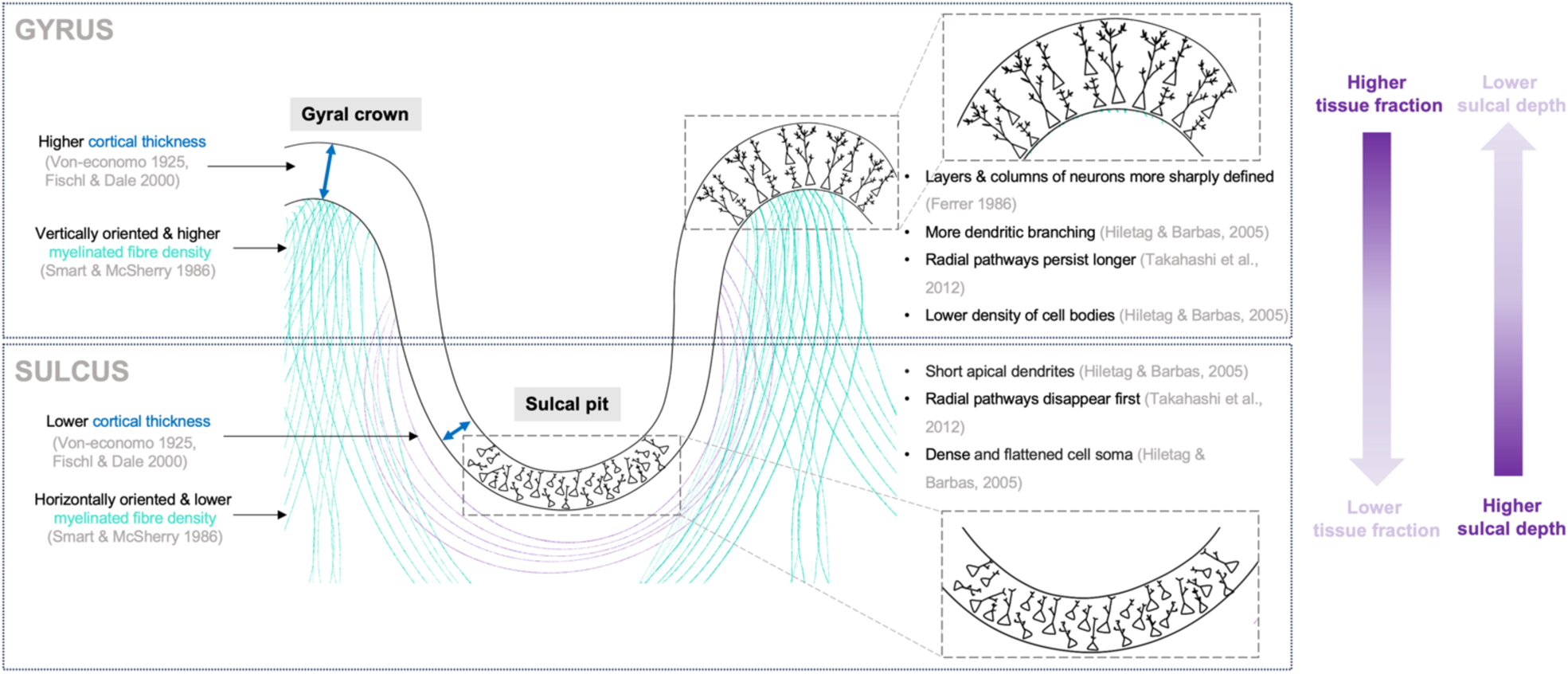
The inverse relationship between tissue fraction and sulcal depth recapitulates adult cytoarchitectural organisation in utero.

### Tissue fraction in the CP and SP decreases prior to sulcal formation

We examined whether regional variations in microstructure predate macrostructural formation by using an age-mismatched coupling analysis, to reinforce the hypothesis that there are microstructural features of the CP and SP have mechanistic importance for the formation of cortical folds. We correlated native tissue fraction values in the youngest subjects in ‘unfolded’ brain regions (25 GW – 29 GW) with 35 GW group-average sulcal depth values, effectively back-projecting a map of folded brain regions onto the unfolded brain (Figure 4). We identified significant negative correlations between future sulcal depth and tissue fraction values across several unfolded brain regions in the youngest subjects. With this analysis, we suggest that dynamic changes in CP and SP microstructure may occur immediately prior to the formation of cortical convolutions, by extension implying that the developmental processes captured by the diffusion signal in these regions might play a role in driving the cortical folding process.

### The mechanistic implications of the relationship between tissue fraction and sulcal depth

The late second to third trimester is characterised by synaptogenesis, increasing maturation of dendritic and axonal arbours, extending glial processes, leading to generalized increases in the complexity of the neuropil (Kostović and Jovanov-Milosević, 2006). Our previous work demonstrated that tissue fraction values reflect the dynamic effect of these neurobiological processes on the biophysical properties of different fetal compartments (Wilson *et al.,* 2023). The decreased tissue fraction in regions that are about to fold suggests that the regional variation in the density and complexity of neuropil in the CP and SP might play a mechanistic function in the onset of folding across the brain. Variations in the density and morphology of neurons accumulating in the CP have been observed in previous work and are thought to be driven in part by differential patterning of gene expression in the outer subventricular zone (OSVZ) and ventricular zone (VZ), which also reliably map the prospective location of sulcal and gyral folds (Gray, Leber and Sanes, 1990; De Juan Romero and Borrell, 2015; Llinares-Benadero and Borrell, 2019). Recently published work has explored this at the histological level in primates, and suggested that cortical expansion and subsequent gyrification are driven by rapid, post-neurogenic neuropil growth, addition of glial cells, and expansion of large white matter tracts (Reillo *et al*., 2011; Nonaka-Kinoshita *et al*., 2013; Stahl *et al*., 2013; Shinmyo *et al*., 2022; Rash *et al*., 2023).

### Age-mismatched analysis highlights the insula as an exceptional region, possibly indicative of unique cortical folding mechanisms in this area

Although this inverse relationship between tissue fraction and sulcal depth was present across many neighbourhoods, a notable exception was the insular cortex, where the age-mismatched analysis highlighted positive correlations between tissue fraction and future sulcal depth. This result is congruent with previous work highlighting the divergent growth and folding trajectory of the insula compared to other lobes (Türe *et al.,* 1999; Mallela *et al.,* 2023). The insula has a unique, less convoluted morphology, with shallower and straighter gyri, with evidence demonstrating that it acquires an adult-like gyration and architectonic pattern during the last trimester of gestation (González-Arnay, González-Gómez and Meyer, 2017). The distinctive properties of this region, also highlighted by our analysis, suggest regionally distinct biological processes may be responsible for the global geometry of the adult brain (Mallela *et al.,* 2020). A more thorough investigation of insula maturation in the fetal brain, and the relationship between surface features and the underlying microstructure, would be beneficial to understand the interaction between different mechanisms of gyrification.

## Conclusion

Projecting the SP and CP tissue and fluid fraction to the white matter surface recapitulated the regionally variant microstructure between these fetal compartments described in our previous work (Wilson *et al.,* 2023), and the corresponding non-linear maturational trajectories with GA. In this work, we uncover a direct link between local micro and macrostructure that offers mechanistic insight about how gyrification begins in the fetal brain, with clear implications for understanding how folding abnormalities arise in utero. Our observations show that the regional variations in microstructural maturity are also meaningfully correlated with emerging cortical folding patterns. This may reflect differences in neurogenesis and the tangential dispersion of migrating neurons, leading to different initial densities of accumulated neurons in the CP before folding (Reillo *et al*., 2011; Borrell and Reillo, 2012). Overall, this study supports hypotheses that allude to an important role of neuronal and glial cellular morphological maturation in producing intracortical forces to drive the formation of primary folds (Richman *et al*., 1975; Wang *et al*., 2017; Llinares-Benadero and Borrell, 2019). Future work would benefit from acquiring longitudinal data at specific time points across the fetal period to test this relationship, building towards establishing causality and clarity about the mechanisms driving gyrification.

## Methods

### Acquisition and pre-processing

112 in utero fetal T2 and diffusion MRI datasets were acquired on a Philips Achieva 3T system with a 32-channel cardiac coil (Price *et al.,* 2019). T2 weighted images were acquired with multiple single-shot turbo spin echo sequences with TE = 250ms, TR = 2265ms (Price *et al.,* 2019). DWI was acquired with a combined spin and field echo (SAFE) sequence at 2 mm isotropic resolution and multi-shell high angular resolution diffusion encoding (15 volumes at b=0s/mm², 46 volumes at b=400s/mm², and 80 volumes at b=1000s/mm²) (Christiaens *et al*., 2019). T2 datasets were reconstructed to 0.5mm isotropic resolution using an automated bespoke pipeline (Cordero-Grande *et al*., 2019). HARDI datasets were reconstructed to 0.8mm, using a data driven representation of the spherical harmonics and radial decomposition (SHARD). The SHARD pipeline caters to the motion corrupted fetal data, using dynamic distortion correction and slice-to-volume motion correction framework (Cordero-Grande *et al*., 2019; Christiaens *et al*., 2021).

### Image registration & segmentation

Alignment between diffusion and T2w volumes for each subject used FLIRT boundary-based registration in FSL (Jenkinson and Smith, 2001; Jenkinson *et al*., 2012). For registration of individual subject T2w images to a common space, a spatiotemporal atlas of the fetal brain was used (https://gin.g-node.org/kcl_cdb/fetal_brain_mri_atlas) (Uus et al., 2023), constructed using the Medical Image Registration ToolKit (MIRTK) atlas generation pipeline (Schuh et al., 2018) (https://biomedia.doc.ic.ac.uk/software/mirtk/). Cortex, white matter and CSF probability maps were generated using the Developing brain Region Annotation With Expectation-Maximization (Draw-EM) module of MIRTK. An automated deep learning pipeline was then used to segment T2w brain images into different tissue types, including deep grey matter, white matter, CSF and cortex (Makropoulos *et al*., 2014). A Convoluted Neural Network (CNN) was trained on manually refined labels propagated from the fetal dHCP atlas (Makropoulos *et al*., 2018; Schuh *et al*., 2018). For each subject, a combination of the T2w volume and the cortex probability map was used for multi-channel ANTs non-linear symmetric diffeomorphic image registration, creating warps between native T2 space and age-matched weeks of the atlas. The multi-channel approach improved accuracy at cortical boundaries.

### Surface reconstruction, registration, and vertex correspondence

To extract hemispheric surfaces on the inner cortical plate boundary, we applied the marching-cube algorithm in CIVET-2.1.0 software package (Lepage *et al*., 2021). Inner parts of the segmented CP were binarized and the initial meshes were tessellated by fitting the boundary of the inner part of CP. We resampled the initial meshes to the standard format surfaces containing 81,920 triangles and 40,962 vertices (Lepage *et al*., 2021). To eliminate small geometrical noise, Taubin smoothing was applied to the surfaces (Taubin, 1995). Then, the surfaces were registered to a 28 GW template surface to guarantee the vertex correspondence among individual surfaces. We used a 2D sphere-to-sphere warping method, which searches optimal correspondence of vertices based on folding similarity between individual and template surfaces (Robbins, 2004; Boucher, Whitesides and Evans, 2009).

### Surface metric calculation

To describe the formation of sulci, we chose sulcal depth, which was calculated using the adaptive distance transform (ADT) method (Yun *et al*., 2013). This metric was selected because it has an intuitive meaning that can be easily interpreted for the purposes of local coupling analysis.

### Diffusion modelling: using DTI and MSMT-CSD to derive maps of diffusion metrics

The approach to modelling the DWI signal is described in previous publication (Wilson *et al.,* 2023). Briefly, white matter response functions were extracted from areas of relatively mature white matter (corticospinal tract and corpus callosum) using the Tournier algorithm (Tournier, Calamante and Connelly, 2013; Tournier *et al*., 2019). CSF responses were extracted using masks of the ventricles using the Dhollander algorithm in MRtrix3 (Jeurissen *et al*., 2014; Tournier *et al*., 2019). To obtain group-average response functions, the white matter response functions of the oldest 20 subjects were averaged (approximating relatively mature anisotropic fetal white matter) and the CSF (isotropic) group-average response function was calculated from the whole cohort. Using these two average response functions, the dMRI signal of all subjects was subsequently decomposed into a tissue and fluid component, using the tournier algorithm applied via the MRtrix3 software package (Tournier *et al*., 2019), and resulting components were intensity normalised for each subject (Raffelt *et al*., 2011). The tissue and fluid component maps were normalised to 1, for ease of interpretation, then labelled as tissue and fluid ‘fraction’.

### Projecting from the surface boundary into diffusion maps

The inverse transform was applied to the surfaces to register them back to native T2 space, and surface coordinates were converted to cartesian space (x,y,z). This preserved the vertex correspondence between all subjects and the 28 GW surface template, such that the surface normal could be used to extrapolate from the white matter boundary into the T2-aligned diffusion maps at equivalent points across the surface. To obtain estimates of microstructure in the SP and CP, we projected ±2mm (twice the surface normal) inside (SP) and outside (CP) the white matter boundary, interpolating into volumetric space to create a ribbon of voxels that was used to sample the maps of diffusion metrics.

### Coupling analysis within-subject

To understand the local relationships within cortical areas of individual subjects, correlation analysis was performed within vertex neighbourhoods, to calculate the ‘coupling’ between microstructure and macrostructure (adapted from Vandekar *et al*., 2016). Custom-built MATLAB analysis pipelines were written, to find neighbours around a central vertex up to 5 degrees of separation on the surface mesh. To determine the appropriate neighbourhood size, we established a compromise that allowed us to include a sufficient number of vertices to correlate microstructure and surface metrics (∼150 vertices per patch), but would also subdivide the primary sulci into multiple patches.

To describe the relationship between diffusion and surface metrics in each neighbourhood, we used the Pearson correlation coefficient (r) and corresponding p value was calculated. The subjects were grouped into bi-weekly bins, and a one sample t-test was used on the Pearson r values at each vertex, within each age group, to identify significant vertices. To correct for multiple comparisons across all vertices, we controlled for the false discovery rate (FDR) (Benjamini and Hochberg, 1995), identifying significant features while incurring a relatively low proportion of false positives. The mean r value is displayed at the vertices which survived FDR correction in each age bin (23, 25, 27.. 35 GW).

### Age-mismatched coupling analysis

Using surface registration to obtain vertex correspondence between subjects and a central timepoint template (28 GW) across the whole surface, it was possible to back-project a map of folded brain regions from the oldest fetal timepoint, onto the unfolded brain. The same technique of analysing coupling in local patches was repeated, but instead of correlating native sulcal depth and diffusion metrics, the average sulcal depth values at 35 GW were correlated with native diffusion metrics (tissue and fluid fraction). To display the results across the whole cohort, the r values were averaged between subjects in biweekly bins. Statistical testing for significant correlations was repeated as above, correcting for FDR across all vertices, then displaying significant r values.

## Ackowledgements and Funding

We thank the patients who agreed to participate in this work and the staff of St Thomas’ Hospital London. This work was supported by the European Research Council under the European Union Seventh Framework Programme (FP/2007–2013)/ERC Grant Agreement No. 319456. We acknowledge infrastructure support from the National Institute for Health Research (NIHR) Mental Health Biomedical Research Centre (BRC) at South London and Maudsley NHS Foundation Trust, King’s College London, and the NIHR-BRC at Guy’s and St Thomas’ NHS Foundation Trust. We also acknowledge grant support in part from the Wellcome Engineering and Physical Sciences Research Council (EPSRC) Centre for Medical Engineering at King’s College London (WT 203148/Z/16/Z) and the Medical Research Council (UK) (MR/K006355/1) and (MR/L011530/1). SW was supported by PhD funding from the UK Medical Research Council/Sackler Foundation (MR/P502108/1). JO is supported by a Sir Henry Dale Fellowship jointly funded by the Wellcome Trust and the Royal Society (206675/Z/17/Z). JO, MR, ADE, JO, and TA received support from the Medical Research Council Centre for Neurodevelopmental Disorders, King’s College London (MR/N026063/1). TA was supported by an MRC Clinician Scientist Fellowship (MR/P008712/1). VK and TA were supported by an MRC Transition Support Award (MR/V036874/1). TA received support from the Medical Research Council Centre for Neurodevelopmental Disorders, King’s College London [MR/N026063/1]. Support for this work was also provided by the NIHR-BRC at Kings College London, Guy’s and St Thomas’ NHS Foundation Trust in partnership with King’s College London, and King’s College Hospital NHS Foundation Trust. KI, EG and HJY are supported by the National Institute of Neurological Disorders and Stroke (R01NS114087) and National Institute of Biomedical Imaging and Bioengineering (R01EB031170) of the National Institutes of Health (NIH).

## Notes

### Competing Interest Statement

The authors have declared no competing interest.

### Summary of Updates

The manuscript has been reformatted with some revised interpretation and discussion, but the major findings of the article remain the same.

