## Supplementary information for "Dynamic changes in subplate and cortical plate microstructure at the onset of cortical folding in vivo"

**Cohort**

Participants were recruited and scanned according to dHCP protocol, which was approved by the UK Health Research Authority (Research Ethics Committee reference number: 14/LO/1169). Written parental consent was obtained in every case for imaging and open data release of the anonymised data. All data was acquired in St Thomas Hospital, London, United Kingdom. GA was determined by sonography at 12 post-ovulatory weeks as part of routine clinical care. Consenting mothers were scanned between 24 and 38 GW with a Philips Achieva 3T system, with a 32-channel cardiac coil in maternal supine position.

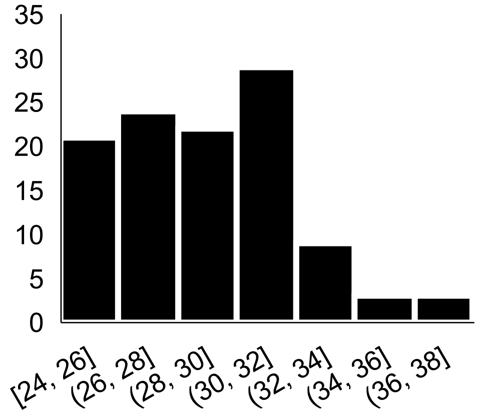


**Supplementary Figure 1.** Gestational Age (GA) distribution of fetal cohort (n = 112 subjects, 68 male and 44 female)

**dMRI acquisition and processing**

dMRI data was collected with a combined spin echo and field echo (SAFE) sequence (Hutter et al., 2018a, Cordero-Grande et al., 2018) at 2 mm isotropic resolution, using a multi-shell diffusion encoding that consists of 15 volumes at b=0 s/mm2, 46 volumes at b=400 s/mm2, and 80 volumes at b=1000 s/mm2 lasting 14 min (Christiaens et al., 2019). For more insight on the choice of b-shell values for this acquisition, readers are directed to Tournier et al., 2019. The protocol also included the collection of structural T2w, T1w, and fMRI data, for a total imaging time of approximately 45 min (Price et al., 2019).

dMRI data were processed using a bespoke pipeline (Christiaens et al., 2019a) that includes generalised singular value shrinkage image denoising and debiasing from complex data (Cordero-Grande et al., 2019), dynamic distortion correction of susceptibility-induced B0 field changes using the SAFE information (Ghiglia and Romero, 1994; Cordero-Grande et al., 2018; Hutter et al., 2018b) and slice-to-volume motion correction based on a multi-shell spherical harmonics and radial decomposition (SHARD) representation (Christiaens et al., 2021).

**Quality control**

Quality control (QC) was implemented using summary metrics based on the gradient of the motion parameters over time and the percentage of slice dropouts in the data (Christiaens et al., 2021, Wilson et al., 2023). This was followed up with expert visual assessment, which considered any residual or uncorrected artefacts. Image sharpness, residual distortion, and motion artefacts were visually assessed and scored between 0 and 3, with 0 (=failure, e.g. because the subject moved out of the field of view) to 3 (=high quality), based on the mean b=0, b=400, and b=1000 images and the ODFs estimated with MSMT-CSD. See Appendix 1—figure 2 for examples of subjects that were excluded.

After co-registration, the QC scores were checked and validated again by different authors (DC, AU, SW), only subjects scoring 2 or 3 that were well aligned in T2 space were admitted to this study. Based on the above criteria, 140 of the 300 subjects that were pre-processed were classified as high-quality reconstructions for both DWI and T2 modalities.

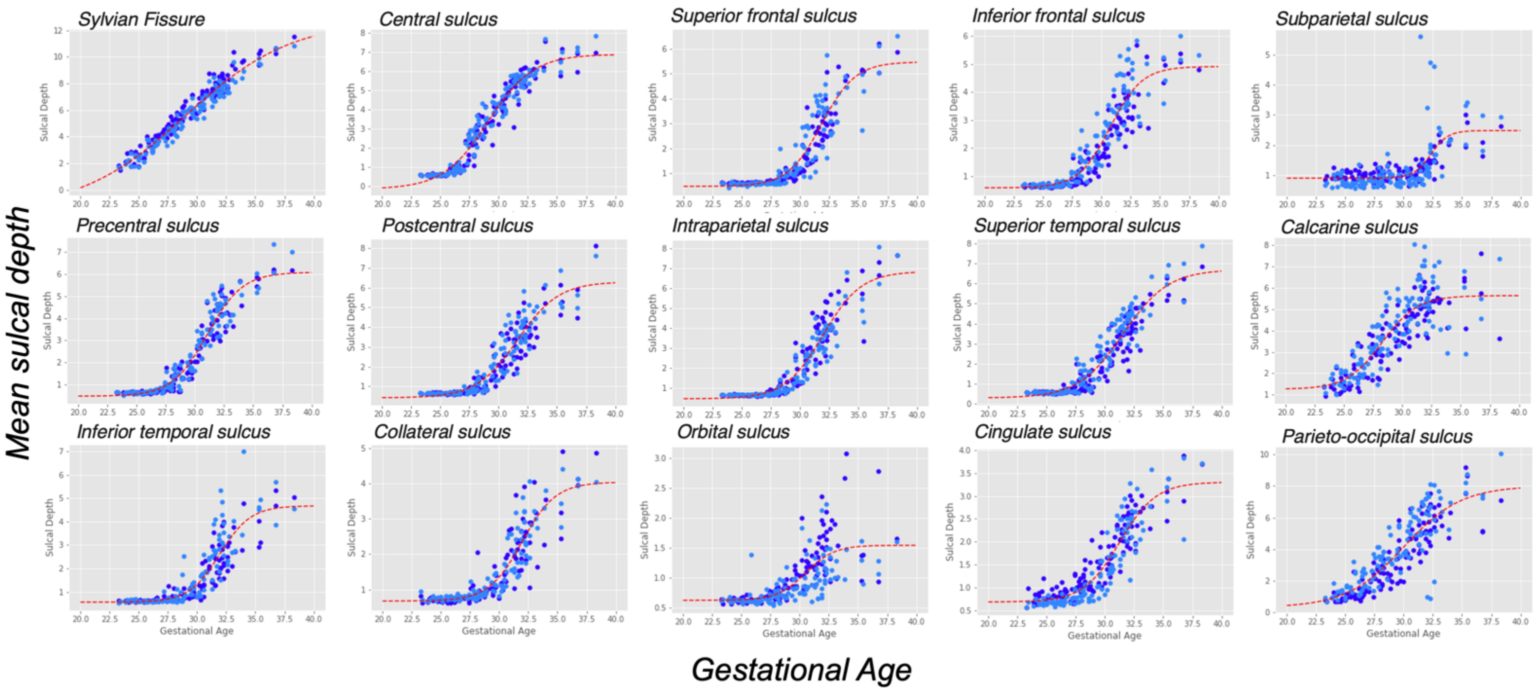


**Supplementary Figure 2.** Mean sulcal depth in each subject in 15 major primary sulci, for the left (light blue) and right (dark blue) hemispheres. Each curve was fit with a sigmoid to describe the growth and formation of the sulci.

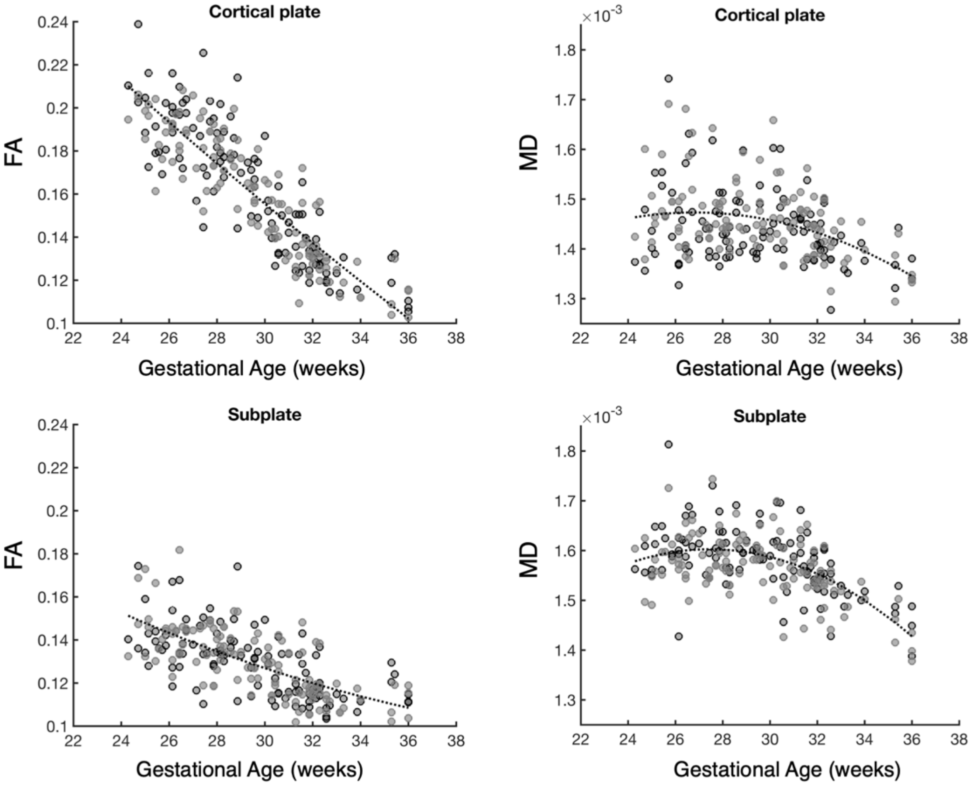


**Supplementary Figure 3.** Mean Diffusion Tensor Metrics (Fractional Anisotropy (FA)) and Mean Diffusivity (MD) in the Cortical Plate and Subplate, charted against gestational age.
